# An Open-Source Deep Learning-Based Toolbox for Automated Auditory Brainstem Response Analyses (ABRA)

**DOI:** 10.1101/2024.06.20.599815

**Authors:** Abhijeeth Erra, Jeffrey Chen, Cayla M. Miller, Elena Chrysostomou, Shannon Barret, Yasmin M. Kassim, Rick A. Friedman, Amanda Lauer, Federico Ceriani, Walter Marcotti, Cody Carroll, Uri Manor

**Affiliations:** Data Institute, University of San Francisco, San Francisco, CA; Dept. of Cell & Developmental Biology, University of California San Diego, La Jolla, CA; Dept. of Otolaryngology, University of California San Diego, La Jolla, CA; Depts. of Otolaryngology-Head and Neck Surgery and Neuroscience and Center for Functional Anatomy and Evolution, Johns Hopkins University School of Medicine, Baltimore, MD; School of Biosciences, University of Sheffield, Sheffield, S10 2TN, UK; Neuroscience Institute, University of Sheffield, Sheffield, S10 2TN, UK; Dept. of Mathematics and Statistics, University of San Francisco, San Francisco, CA; Halıcıoğlu Data Science Institute, University of California San Diego, La Jolla, CA

## Abstract

Hearing loss is a pervasive global health challenge with profound impacts on communication, cognitive function, and quality of life. Recent studies have established age-related hearing loss as a significant risk factor for dementia, highlighting the importance of hearing loss research. Auditory brainstem responses (ABRs), which are electrophysiological recordings of acoustically evoked synchronized neural activity from the auditory nerve and brainstem, serve as *in vivo* correlates for sensory hair cell and synaptic function, hearing sensitivity, and other critical readouts of auditory pathway physiology, making them highly valuable for both basic neuroscience and clinical research. Despite their utility, traditional ABR analyses rely heavily on subjective manual interpretation, which may introduce variability and pose challenges for reproducibility across studies. Here, we introduce Auditory Brainstem Response Analyzer (ABRA), a novel suite of open-source ABR analysis tools powered by deep learning, which automates and standardizes ABR waveform analysis. ABRA employs convolutional neural networks trained on diverse datasets collected from multiple experimental settings, achieving rapid and unbiased extraction of key ABR metrics, including peak amplitude, latency, and auditory threshold estimates. We demonstrate that ABRA’s deep learning models provide performance comparable to expert human annotators while dramatically reducing analysis time and enhancing reproducibility across datasets from different laboratories. By bridging hearing research, sensory neuroscience, and advanced computational techniques, ABRA facilitates broader interdisciplinary insights into auditory function. An online version of the tool is available for use at no cost at https://abra.ucsd.edu.

## Introduction

Hearing loss is a prevalent and debilitating condition affecting hundreds of millions worldwide. In addition to significantly diminishing quality of life, age-related hearing loss has emerged as a major risk factor for cognitive decline and dementia, underscoring the urgent need for better research tools and treatment strategies. Both age-related hearing loss and dementia involve the progressive loss of synapses (synaptopathy) - in the cochlea and brain, respectively - highlighting common neurodegenerative mechanisms that warrant intensive study (O’Leary et al. 2017; Bramhall et al. 2018; Loughrey et al. 2018; Ray et al. 2018; Liu et al. 2020; Livingston et al. 2024; Park et al. 2024).

Among the most powerful approaches to assess auditory function are auditory brainstem response (ABR) recordings, which objectively measure electrical activity along the auditory neural pathway, from cochlear inner hair cells through the brainstem (Eggermont 2019; Kim et al. 2022; Burkard and Don 2012; Ingham et al. 2011; Møller and Jannetta 1985; Xie et al. 2018). In mice, ABRs consist of five characteristic peaks approximately corresponding to neural signals propagating through sequential auditory structures, though some centrally generated waves may reflect concurrent activity in multiple structures (**Figure 1**, Rüttiger et al. 2017; Melcher et al. 1996; Henry 1979; Land et al. 2016). Physiological and mathematical models of ABR morphology have also been developed (Kamerer et al. 2020).

**Figure 1:**
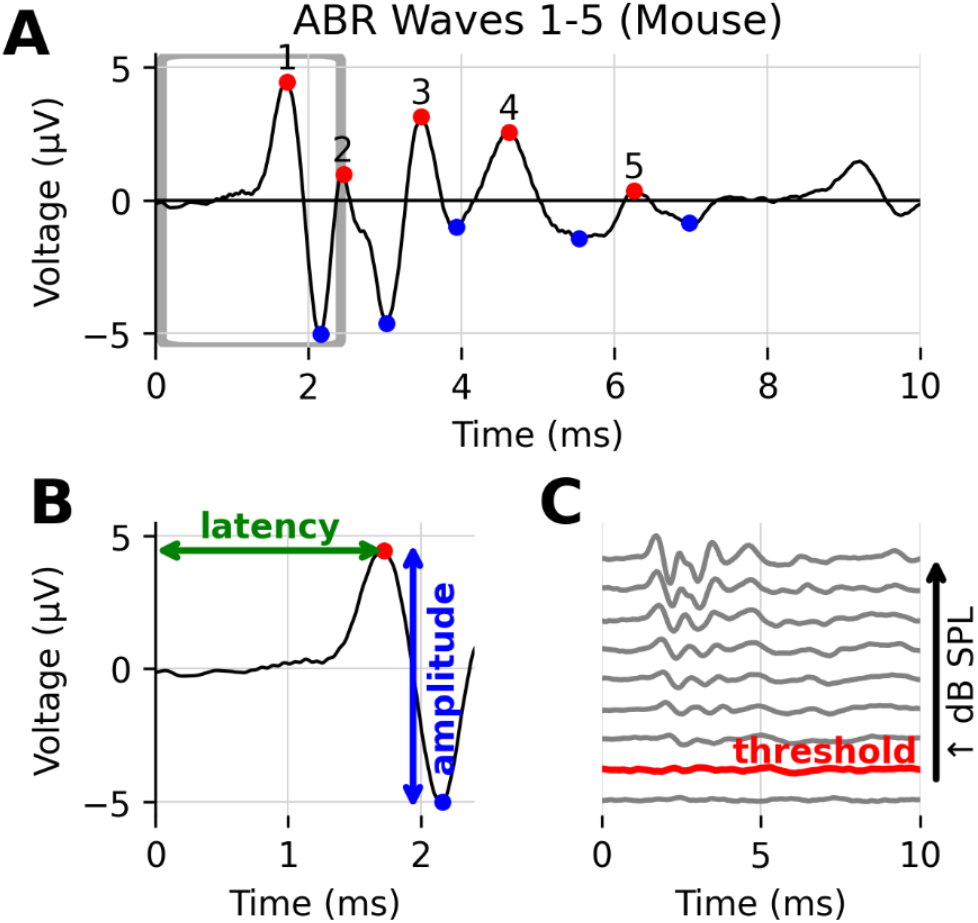
Example of ABR waveforms recorded from a mouse, showing its characteristic features. (A) One 10 ms ABR recording, with the five characteristic peaks denoted by red dots, and their corresponding troughs with blue dots. (B) A close up of the boxed region in (A), showing how latency (time to peak) and amplitude (peak to trough height) are defined for wave 1. (C) Several ABR recordings at varying sound levels, for the same mouse and frequency. The threshold level above which the ABR response is indicative of hearing is shown in red.

Two particularly informative measurements obtained from ABRs—hearing threshold sensitivity and wave 1 characteristics—have been shown to correlate strongly with cochlear-based hearing impairment; specifically, hearing thresholds increase and wave 1 amplitudes are dampened if hair cell synapses are damaged. (Fernandez et al., 2015; Bramhall et al. 2018; Bao et al. 2022; Young, Cornejo, and Spinner 2023). However, current ABR analysis methods typically rely on manual waveform interpretation, which can be subjective, labor-intensive, and prone to inconsistency between individual researchers or labs (Suthakar and Liberman 2019, Schrode et al. 2022).

Heuristic and machine learning computational approaches have been explored for automated ABR analysis (Shaheen et al. 2025). Early methods focused on hand-engineered features and statistical classifiers, such as support vector machines for threshold detection (Acir & Özdamar 2006) or model-based approaches for identifying near-threshold responses (Sánchez et al. 1995). Supervised learning models (i.e. models which learn from data with ground truth labels) like convolutional neural networks (CNNs), gradient boosting machines, and others have been used to accurately analyze suprathreshold ABR waveforms (Wimalarathna et al. 2021, McKearney and MacKinnon 2019) and to assess the degree of synaptopathy in humans (Buran et al. 2022).

In this paper, we introduce the Auditory Brainstem Response Analyzer (ABRA), a collection of novel open-source tools, including machine learning models trained on a diverse set of mouse ABR to enable comprehensive and maximally generalizable mouse ABR analysis. This suite of algorithms automatically detect peaks, estimate thresholds, and quantify latencies. To make these methods broadly accessible, we have packaged them into a user-friendly, browser-based application that also supports batch data import/export, waveform visualization, and interactive 2D/3D plotting. By integrating these diverse functionalities into a unified platform, ABRA aims to streamline ABR data processing and analysis, reduce manual labor, and facilitate standardization and reproducibility across labs. We demonstrate the flexibility and generalizability of these algorithms by benchmarking the performance on ABR datasets collected from three different hearing research labs using distinct experimental protocols and recording settings.

## Methods

### Data Collection

To test the generalizability and flexibility of developed open-source ABR software, we used three distinct datasets from different laboratories to train and evaluate ABRA’s models (**Table 1**). While the three laboratories used a similar overarching methodology, including anesthesia, electrode placement, and range of stimulus sound levels, each used unique experimental protocols, including varying collection software, sound source and stimulus frequencies, and mouse strains, including mouse models of accelerated aging and mice exposed to temporary threshold shift-inducing noise. These differences underscore the flexibility of ABRA in accommodating diverse experimental setups and protocols. Further details on data collection conditions are available in the Supplementary Information (**Supplementary Table S1**).

**Table 1:** Breakdown of mouse and ABR waveform data used from the three different laboratories (Lab A, Lab B, Lab C) in the model split into a train and test datasets. Figures and Tables relevant to a given dataset are enumerated in brackets in the last row.

| Lab/Model | Peak Detection |  | Automatic Thresholding |  |  |
| --- | --- | --- | --- | --- | --- |
|  | Training Data | Test Data | Training Data | Test Data | ABRA vs. EPL-ABR |
| **Lab A** | 32 mice<br>(288 ABRs) | 8 mice<br>(80 ABRs) | 29 mice<br>(5,335 ABRs) | 9 mice<br>(1,393 ABRs) | – |
| **Lab B** | 83 mice<br>(8,635 ABRs) | 21 mice<br>(2,247 ABRs) | 75 mice<br>(11,700 ABRs) | 19 mice<br>(2,966 ABRs) | – |
| **Lab C** | – | – | 10 mice<br>(243 ABRs) | 2 mice<br>(122 ABRs) | 2 mice<br>(122 ABRs) |
| **Total**<br>**[Relevant**<br>**Figures &**<br>**Tables]** | **115 mice**<br>**(8,923 ABRs)**<br>**[Fig. 2]** | **29 mice**<br>**(2,327 ABRs)**<br>**[Fig. 4, 5; Table 2]** | **114 mice**<br>**(17,278 ABRs)**<br>**[Fig. 3]** | **30 mice**<br>**(4,481 ABRs)**<br>**[Figs. 6-8;**<br>**Tables 3, 4]** | **2 mice**<br>**(122 ABRs)**<br>**[Table 5]** |

### Peak Detection

The ABRA toolbox incorporates a two-step peak finding algorithm that leverages Pytorch’s deep learning library and the Scikit-learn library. The first step involves deploying a Convolutional Neural Network (CNN) to predict the location of the wave 1 peak. The CNN was trained on 7,209 ABRs (222 ABRs from 25 mice (Lab A); 6,987 ABRs from 67 mice (Lab B)) labeled with ground truth peak 1 annotations and validated on an additional 1,714 ABRs (66 ABRs from 7 mice (Lab A); 1,648 ABRs from 16 mice (Lab B)). ABR data and analysis of wave 1 from Lab B was previously described (Ceriani et al., 2025). Each training sample’s loss is weighted to ensure that both lab’s data is represented equally in the model training.

Before training the CNN, each ABR waveform was rescaled as follows: a standardized scaler applied over the waveform generated z-scores for each timepoint, and a min-max scaler on those z-scores scaled each wave between 0 and 1. Each input ABR is given as a vector of length 244, covering 10 ms of recording. ABRs not initially sampled at 244 samples per 10 ms were truncated to 10 ms and either downsampled to 244 points using linear interpolation or upsampled to 244 points using cubic spline interpolation. For the downsampled waveforms, we computed the power spectrum of the waveform to ensure the results were not affected by aliasing; we found on average 0.02% and at max 0.75% of the power was above the new Nyquist limit, making aliasing effects negligible. The dataset was split into training, validation, and test sets, with 65%, 15%, and 20% of subjects from each lab going into each set, respectively.

The CNN optimizes squared error loss (L^2^) for the regression task which returns a prediction for the wave 1 peak index. Model architecture and training hyperparameters were determined by a randomized search over convolutional filter sizes, kernel sizes, dropout rates, pooling strategies, learning rates, weight decay values, and early stopping criteria. This approach allowed us to explore a broad space of architectures and select a configuration that provided strong predictive accuracy while maintaining generalization. The validation set was evaluated at each training epoch, and if the validation loss did not decrease over 25 consecutive epochs, training was halted to prevent overfitting. With this model, training stopped after 62 epochs. A simplified representation of the network architecture is shown in **Figure 2**. Model hyperparameters chosen by cross-validation are displayed in **Supplementary Table S2**.

**Figure 2:**
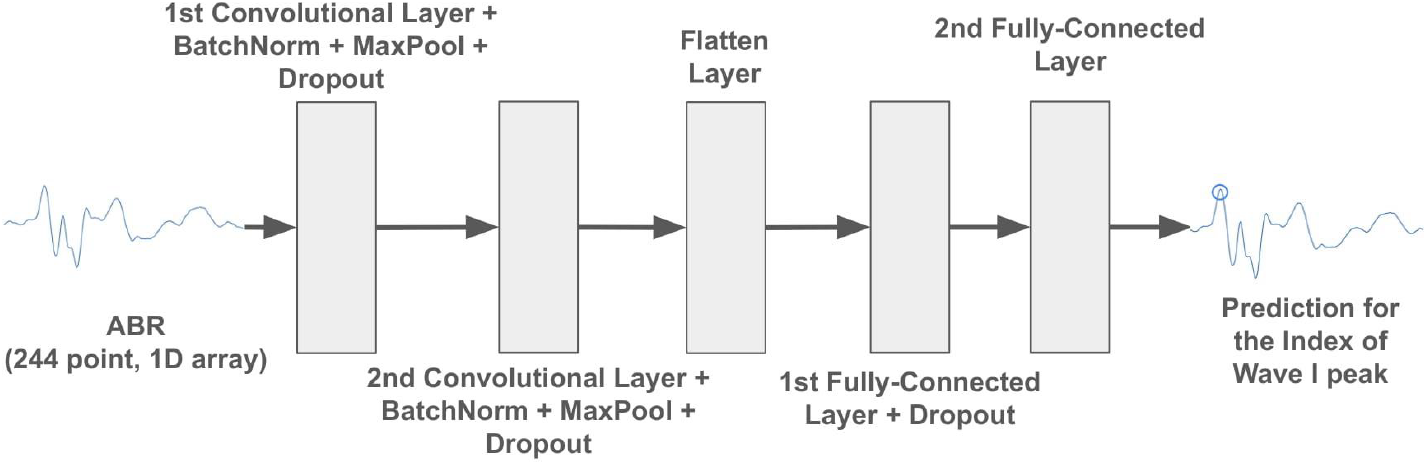
Model architecture for the wave 1 peak finding algorithm. The input (a 244-point ABR waveform spanning 10 ms) passes through two layers of convolution, batch normalization, max pooling (after ReLU activation), and dropout. The dimensionality of the output is then reduced through two consecutive fully-connected layers using ReLU activation which returns the prediction of the time point of the wave 1 peak.

The CNN’s prediction of the wave 1 peak location is often close to, but not exactly at the local peak. With an additional fine-tuning step, this estimate is aligned to the peak location with greater accuracy (**Supplementary Figure S2**). First, the ABR is smoothed using Gaussian smoothing to attenuate or remove nuisance noise to identify peak indices. Then the *find_peaks* method from Scikit-learn is used to identify all candidate peak and trough locations. The best candidate peak 1 is chosen within a ±0.25 ms window from the predicted peak location. If no candidate peaks are found within the window, the window is widened by an additional 0.25 ms until a peak is found. The greatest 4 peaks after this peak are then chosen as peaks 2-5, and troughs chosen which lie between these. The amplitudes are quantified from the original (unsmoothed) waveforms. The parameters for these methods were optimized using ground truth wave 1 latency and wave 1 amplitude for the validation set (1,714 ABRs) and include the following:

a. Window size for the start point for the smoothed waveform being inputted into the *find_peaks* function: 0.41 ms before the CNN prediction for peak 1
b. Minimum allowed time between peaks: 0.66 ms
c. Gaussian smoothing step: *σ*= 1.0
d. Minimum time between candidate troughs: 0.29 ms

Note that these values correspond to integer multiples of the sampling interval (0.041 ms). For example, the “time between troughs” value of 0.29 ms corresponds to 7 sample points. These choices of tuning parameters ensured that the ABRA peak detection method was robust across both high- and low-SNR conditions of ABRs in the training and testing data. Note that this approach may fail for certain conditions where latency is particularly abnormal.

### Supervised Threshold Estimation

The ABRA threshold estimation method uses a binary machine learning classifier to identify individual ABR waveforms as either above or below threshold, where above threshold is taken as the positive class. Once individual waveforms are classified, the hearing threshold for a given frequency is determined as the quietest stimulus level (in dB SPL) for which a subject’s ABR waveform is classified as a hearing response (i.e. above threshold). Three candidate supervised binary classifiers were trained and evaluated: A CNN, an XGBoost classifier, and a Logistic Regression classifier.

Logistic Regression was selected as a baseline linear model because it allowed us to test whether simple linear combinations of waveform amplitudes across timepoints could be sufficient to classify the presence of a signal in the ABR. Its coefficients can provide direct information about which timepoints (i.e. regions around canonical peaks and troughs) contribute most strongly to classification, allowing for interpretability. XGBoost was chosen as a more complex model that could capture higher-order interactions between waveform features. It is known for its strong performance on tabular data. Together, these models provide a spectrum of algorithm complexity against which the CNN’s performance can be compared.

Evaluation metrics include accuracy, true positive rate (TPR; i.e. the proportion of actual above threshold instances that are correctly identified as above threshold), false positive rate (FPR; i.e. the proportion of actual below threshold instances that are incorrectly classified as above threshold), the area under the receiver operator curve (AUCROC) and the area under the precision-recall curve (AUCPR).

The dataset comprised of 21,768 ABR waves from 144 mice (Lab A = 48 mice; Lab B = 104 mice; Lab C = 36 mice), with each wave characterized by its frequency, decibel level, and amplitudes at 244 uniformly distributed sampling points over a 10 ms time window. As for the peak finding model, ABRs not initially sampled at 244 samples per 10 ms were truncated to 10 ms and either downsampled to 244 points with linear interpolation or upsampled to 244 points with cubic spline interpolation. The ABRs were grouped by subject and frequency, then 80% of these waveform stacks were randomly allocated for training and the remaining 20% were designated for testing (see **Table 1** for ABR counts from each lab). This method ensures a representative distribution of ABRs from various subjects and frequencies across the training and testing sets. Accordingly, the training input matrix for the XGBoost classifier and the Logistic Regression Classifier had dimensions of 17,350 x 244, where 17,350 is the total number of training samples and 244 is the number of features, including 244 voltage readings for each ABR.

For the Logistic Regression Classifier and XGBoost Classifier, time warping was used on the ABR trajectories as an additional preprocessing step to align waveform features such as peaks and troughs (see Supplementary section: **ABR Curve Alignment with Time Warping**). ABRs were preprocessed as follows: a standardized scaler applied over the entire stack generated z-scores for each timepoint, and a min-max scaler on those z-scores scaled all values between 0 and 1. To improve generalization, data augmentation techniques were used to increase the sample space used for training. This included noise injection, elastic augmentation through cubic spline interpolation, and time shifting. Thus, the final training input matrix for the CNN had dimensions of 34,700 x 244, where 34,700 is the total number of training samples and 244 is the number of features, representing the 244 voltage readings for each ABR. Each training sample is weighted in the loss calculation to ensure that each lab’s data is represented equally in the model training. The final architecture of the CNN was selected through a randomized hyperparameter tuning search over convolutional filter sizes, kernel widths, pooling strategies, dropout rates, learning rates, and batch sizes. This search ensured that the final network balanced the ability to capture complex waveform features with the need to generalize. The search converged on the final architecture of the CNN as described in **Figure 3**. Model hyperparameters chosen are displayed in **Supplementary Table S2**.

**Figure 3:**
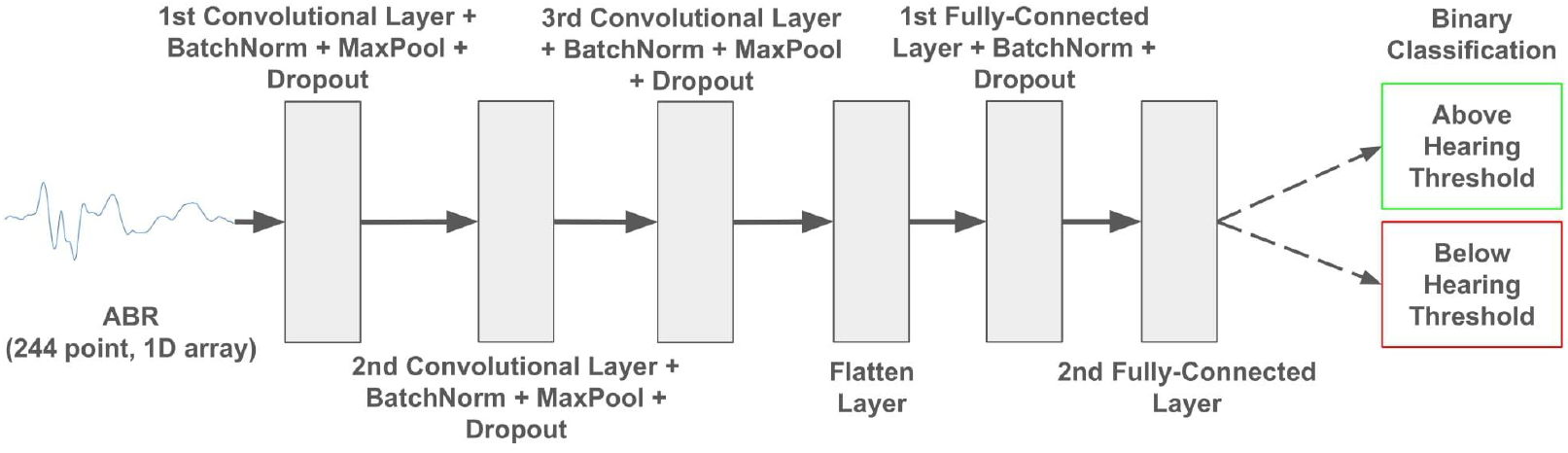
Model architecture for the CNN ABR threshold classifier. The input (a 244-point ABR waveform spanning 10 ms) passes through three sequential layers of convolution, batch normalization, max pooling (after ReLU activation), and dropout. The dimensionality of the output is reduced through two consecutive fully-connected layers using ReLU activation before finally being passed through a sigmoid activation function returning the classification of the ABR.

## Results

### Peak Amplitude and Latency Estimation

To benchmark the performance of the ABRA peak amplitude and latency estimation, we tested its performance on a set of 2,327 ABRs with human-labeled “ground truth” wave 1 amplitude and latency values from Lab A (80 waveforms from 8 mice) and Lab B (2,247 waveforms from 21 mice). The ground truth values for Lab A data were obtained by using visual examination with the BioSigRZ software from Tucker Davis Technologies, while the ground truth values for Lab B data were obtained using a semi-automatic approach using custom software (Ceriani et al. 2025). These ground truth annotations cover a wide range of tested frequencies and therefore latencies (**Supplementary Figure S3**). Though it is possible to make manual adjustments to these predictions, we assess the model here by comparing the automated (i.e. unadjusted) estimates from the ABRA peak finding algorithm vs their corresponding human-labeled ground truth values.

Errors are defined by the differences between the ABRA peak finding algorithm’s automated estimates and the human-annotated ground truth values. The distributions of errors are shown for the latencies and amplitudes of the wave 1 peak in **Figure 4A** and **B**, respectively; summary statistics for errors are reported in **Table 2**. Note that since many measurements had zero error, the average error of -0.004 ms is smaller in magnitude than the measurement precision of 0.041 ms (**Table 2**).

**Table 2:** (related to Figure 5): Table showing the Mean Error Difference and their Standard Errors between ABRA-detected wave 1 latency and amplitude and corresponding ground truth values detected by human reviewers. Two sample t-tests found that mean error differences were not significant for wave 1 latency nor for wave 1 amplitude estimates at the 95% significance level after Bonferroni correction.

|  | Wave 1 Latency | Wave 1 Amplitude |
| --- | --- | --- |
| Mean Difference (± S.E.M.) | Predicted vs. Ground Truth (ms) | Predicted vs. Ground Truth (μV) |
| Lab A ($n_{\text{waveforms}}=80$, $n_{\text{mice}}=8$) | -0.046 ($\pm 0.021$) | -0.115 ($\pm 0.043$) |
| Lab B ($n_{\text{waveforms}}=2,247$, $n_{\text{mice}}=21$) | -0.003 ($\pm 0.002$) | -0.009 ($\pm 0.005$) |
| Overall Test Set | -0.004 ($\pm 0.002$) | -0.013 ($\pm 0.005$) |

**Figure 4:**
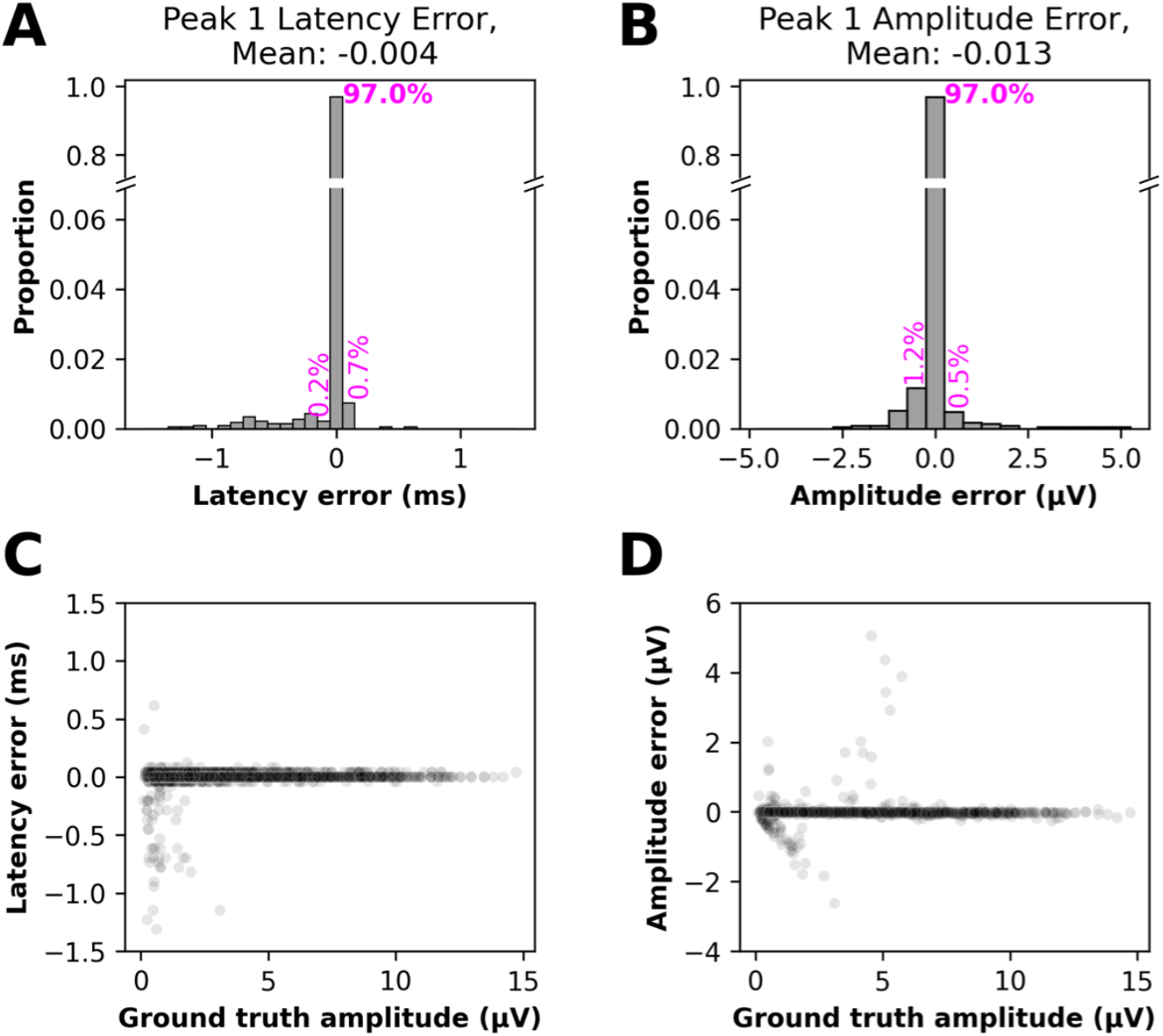
Errors in wave 1 peak quantification vs. ground truth for the ABRA peak finding method. (A) The distribution of errors in the predicted latency of the wave 1 peak (the direct output of the model). 97.0% of errors were within 0.05 ms of the ground truth measurement. (B) The distribution of errors in the wave 1 amplitude (peak to trough), calculated from the predicted wave 1 peak location, together with the estimated trough location. 97.0% of errors were within 0.25 µV of ground truth measurements. (C), (D). Wave 1 latency and amplitude errors vs. ground truth wave 1 amplitude, showing many of the highest errors occur on waveforms with small amplitudes (low SNR). Each predicted peak 1 is shown as a single semi-transparent point. These results are shown by frequency in **Supplementary Figure S4**. n = 2327 ABR waveforms in the test set. Related statistics are listed in **Table 2**.

The root mean squared error of the predicted peak 1 amplitude was *RMSE*_*a*_ *= 0*.*245* µV, *95% CI: 0*.*235-0*.*255* µV. The root mean squared error of the predicted peak 1 latency was *RMSE*_*τ*_ *= 0*.*092 ms, 95% CI: 0*.*088-0*.*095* ms, while the mean absolute latency error (*MAE*_*τ*_ = *0*.*022* ms), or about 0.2% of the total sweep length. These MAE intervals provide assurance that the prediction errors for both amplitude and latency metrics are well bounded. However, the RMSE values are approximately 4x higher than the MAE, indicating that while most errors are small, a few rare instances of larger errors are present, primarily in low SNR conditions (**Figure 4C**). Occasional amplitude errors at moderate amplitudes are often the result of misplaced trough 1 predictions (**Figure 4D, Supplementary Figure S5**).

These comparisons show that the ABRA-generated peak estimates generally agree with human-labeled ground truth latency and amplitude estimates. **Figure 5** displays examples of both correctly identified and erroneous peak 1 predictions, including those at low SNR.

**Figure 5:**
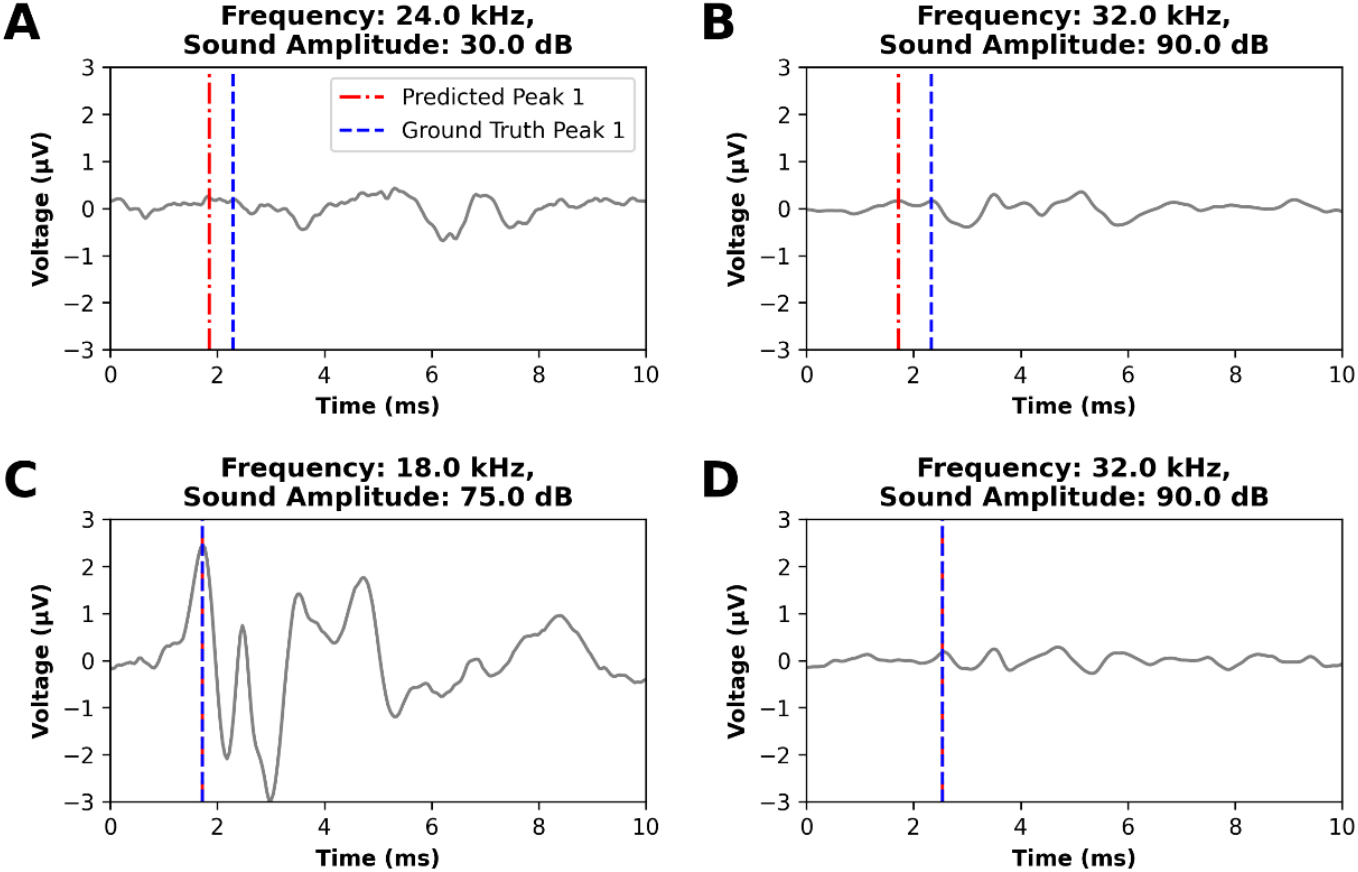
Examples of error and success cases in ABRA automated wave 1 peak detection. (A, B) For ABRs close to the hearing threshold and/or with low SNR, the ABRA peak detection algorithm sometimes identified the incorrect peak. (C) and (D) display examples of ABR waveforms with varying signal to noise ratios for which ABRA matched the ground truth.

### ABR Classification and Threshold Estimation Results

The performance of our ABR classifiers for threshold detection was assessed on the testing set of 4,490 ABR waveforms. Performance metrics are shown in **Figure 6**, and pairwise comparisons for these metrics are provided in **Table 3**. As a simple and interpretable model, logistic regression was used as a baseline for the binary classification task. It achieved an accuracy of 73.01%, a True Positive Rate (TPR), sometimes referred to as recall or sensitivity, of 74.68%, and an Area Under the Receiver Operating Characteristic Curve (AUCROC) of 0.8166. However, its performance was significantly outperformed by both the CNN and XGBoost models. The CNN surpassed XGBoost and the baseline Logistic Regression models in terms of all metrics, indicating the CNN’s enhanced ability to correctly identify ABR thresholds. The CNN model achieved both an AUCROC and an AUCPR of about 0.99 (**Figure 6**). These high values suggest strong overall discrimination at different decision thresholds (AUCROC, **Figure 7**) and excellent performance in handling class imbalance (AUCPR), reflecting robust sensitivity and precision.

**Table 3:** Comparative analysis of performance metrics between the three machine learning models for ABR threshold classification (related to Figures 6, 7). Metrics were calculated on the test set of 4,481 ABR waveforms, and compared between the Convolutional Neural Network (CNN), XGBoost (XGB), and Logistic Regression (LR) models. The CNN model outperforms the XGB model across all metrics. Both CNN and XGB outperform the LR model across all metrics. All statistical tests indicated highly significant differences, with p-values less than 1x10^-16^ (the numerical precision limit for double-precision floating point). Significance was calculated using two-sample proportion tests (Accuracy, TPR, FPR), DeLong’s test (AUCROC), or bootstrapping (AUCPR). A Bonferroni correction was applied to address multiple testing.

| Metric | Comparison | Difference Estimate | 95% CIs for differences |
| --- | --- | --- | --- |
| Accuracy | CNN vs. XGB | 0.0431 | (0.0428, 0.0433) |
|  | CNN vs. LR | 0.2200 | (0.2197, 0.2204) |
|  | XGB vs. LR | 0.1770 | (0.1766, 0.1773) |
| AUCROC<br>Area Under the Receiver Operating<br>Characteristic Curve | CNN vs. XGB | 0.0220 | (0.0174, 0.0266) |
|  | CNN vs. LR | 0.1723 | (0.1580, 0.1866) |
|  | XGB vs. LR | 0.1503 | (0.1365, 0.1641) |
| AUCPR<br>(Area Under the Precision-Recall Curve) | CNN vs. XGB | 0.0104 | (0.0083, 0.0128) |
|  | CNN vs. LR | 0.0824 | (0.0758, 0.0893) |
|  | XGB vs. LR | 0.0720 | (0.0655, 0.0787) |
| TPR<br>(True Positive Rate) | CNN vs. XGB | 0.0361 | (0.0358, 0.0365) |
|  | CNN vs. LR | 0.2141 | (0.2136, 0.2146) |
|  | XGB vs. LR | 0.1780 | (0.1775, 0.1785) |
| **TNR**<br>(True Negative Rate) | CNN vs. XGB | 0.0562 | (0.0554, 0.0570) |
|  | CNN vs. LR | 0.2313 | (0.2302, 0.2323) |
|  | XGB vs. LR | 0.1751 | (0.1740, 0.1762) |

**Figure 6:**
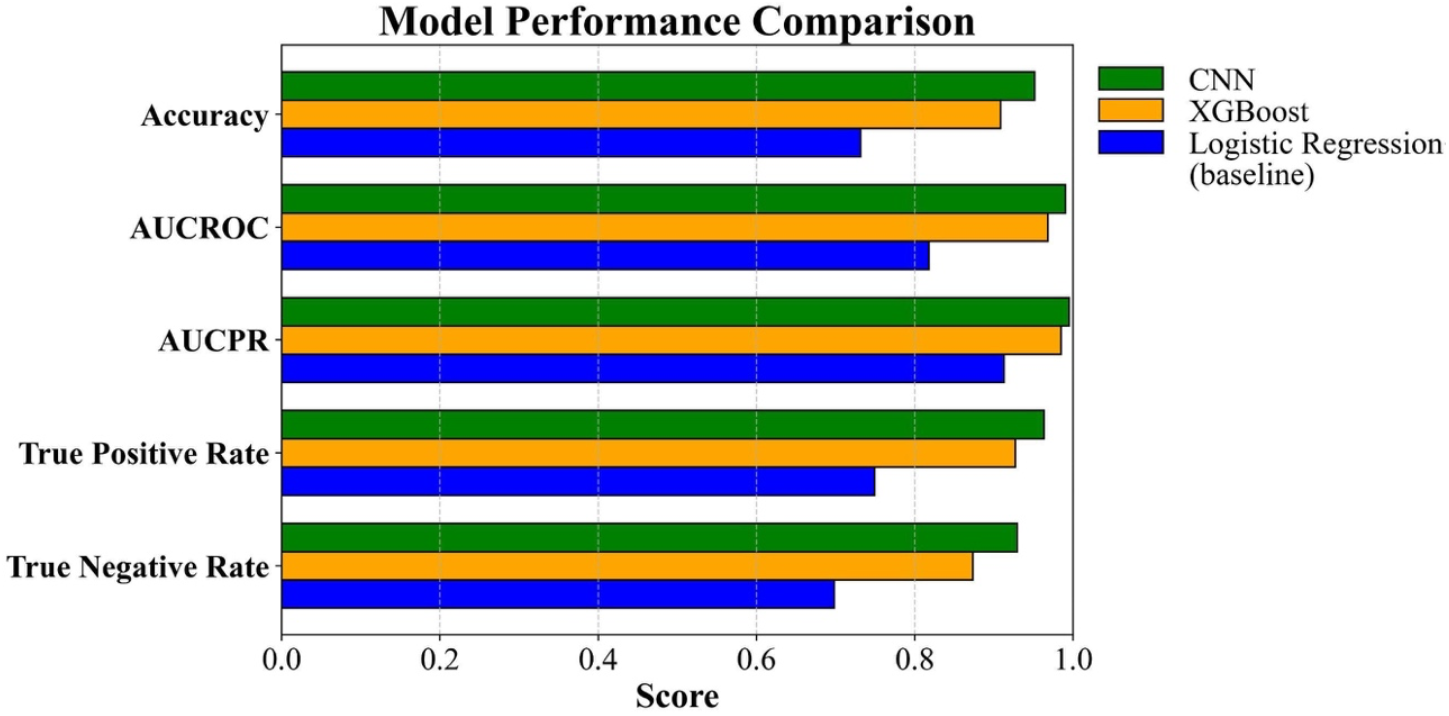
Comparative analysis of machine learning models for ABR threshold classification. Horizontal bar chart illustrating the performance of three machine learning models (top to bottom): Convolutional Neural Network (CNN), XGBoost, and Logistic Regression (baseline). The metrics used for comparison are Accuracy, Area Under the Receiver Operating Characteristic Curve (AUCROC), Area Under the Precision-Recall Curve (AUCPR), True Positive Rate, and False Positive Rate. Note that the false positive rate (FPR) is equal to the complement of the specificity, or true negative rate.

**Figure 7:**
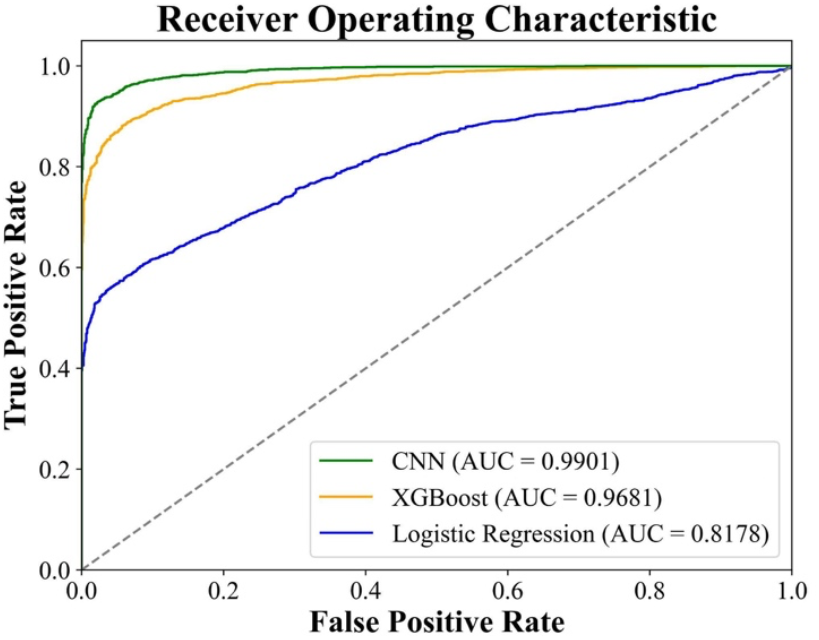
Receiver Operating Characteristic (ROC) curves for the three machine learning models for ABR threshold classification. The ROC curve demonstrates the performance of each ABR classifier (top to bottom: CNN, XGBoost, and Logistic Regression) at all classification thresholds. The area under the ROC curve (Area Under Curve; AUC) represents each ABR classifier’s overall ability to distinguish between above-threshold and below-threshold ABR responses. The ROC curve for the CNN shows slight outperformance of the XGBoost classifier, and both more significantly outperform the Logistic Regression classifier.

The error in our method may partly stem from variations in the subjective determination of thresholds by different researchers, which were used as the ground truth for training our model. To quantify the variability between expert annotators, we sampled 90 ABRs (9 mice at 10 frequencies) from Lab B and asked an expert from Lab A to independently determine the thresholds. The difference between the thresholds assigned by Rater 1 (Lab B) and Rater 2 (Lab A) represents the interrater error. We then trained the CNN on only data from Labs A and C, so that data from Rater 1 was not included in training. On average, the interrater error was comparable to that of the CNN on the sample of Lab B data (**Figure 8**). While the mean absolute errors were similar, we also assessed the accuracy of the CNN predictions within 5 dB and 10 dB envelopes. At the 5 dB threshold, interrater accuracy exceeded that of the CNN by 1-3%, while at the 10 dB threshold, the two were indistinguishable (**Table 4**).

**Table 4:** Inference for differences in accuracy between inter-rater accuracy (IR) and the Convolutional Neural Network (CNN) model in threshold estimation (related to Figure 8). Within a 10 dB SPL envelope, no significant difference between the CNN baseline and inter-rater accuracy was detected, suggesting CNN is performing at a comparable level as a human reviewer at this precision. At the more precise envelope of 5 dB SPL, there was a significant difference of 1-3% after Bonferroni correction. Significance level notation after applying Bonferroni correction: 0.001(***).

| Accuracy Envelope | Comparison | Accuracy Difference | 95% CIs for Accuracy Difference | *p-value* | Significance |
| --- | --- | --- | --- | --- | --- |
| Within 5 dB SPL | CNN vs. IR | -0.0222 | (-0.0310, -0.0134) | $1.5578 \times 10^{-6}$ | *** |
| Within 10 dB SPL | CNN vs. IR | 0.0000 | (-0.0046, 0.0046) | 1.0 | NS |

**Figure 8:**
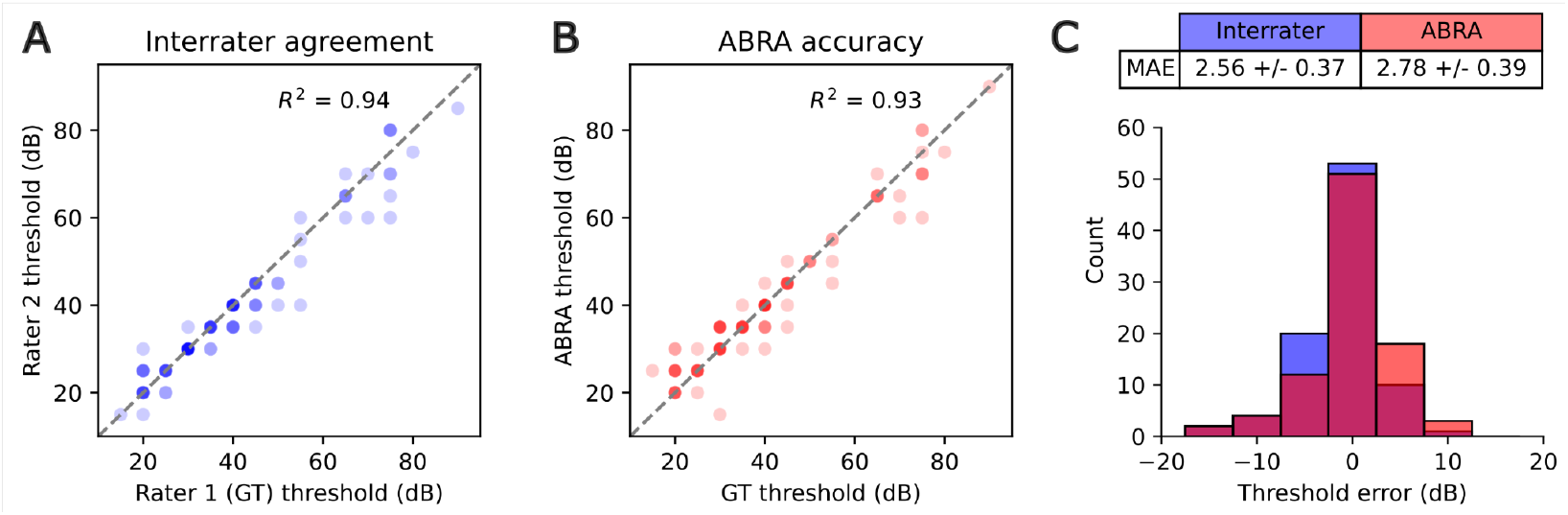
Threshold estimation accuracy of the ABRA thresholding algorithm and agreement among multiple expert raters. (A,B) Comparison of threshold estimates between ground truth (GT) rater and a second expert (A) and between GT and the CNN-based ABRA thresholding method (B), across 90 measurements. Each threshold estimate is displayed as a semitransparent point such that darker points represent multiple overlapping values. (C) Distribution of threshold errors for the same 90 measurements, with interrater differences in blue and ABRA thresholding errors in red. The absolute errors were not statistically different between interrater and ABRA comparisons (p=0.68, t-test). The thresholds were estimated for 9 mice at 10 frequencies from Lab B. All data from Lab B was excluded from training and testing the CNN for this experiment.

The CNN model’s superior performance at the 5 dB and 10 dB envelopes, along with its ability to handle the temporal nature of the data, makes it the optimal choice for this task. Moreover, the CNN achieves similar performance to the inter-rater comparison, indicating that its performance in estimating hearing thresholds is on par with the consensus of two human experts using standard methods. This suggests that the CNN model can function as a reliable tool for estimating hearing thresholds, providing a machine learning-based approach that matches human expert performance.

The performance of our threshold estimation technique was compared against the cross-correlation algorithm embedded in EPL-ABR (Suthakar & Liberman 2019) on a separate dataset of ABR waveforms from Lab C (**Table 5**). This smaller set of ABR waveforms (N = 122) was selected because EPL-ABR’s threshold estimation software requires data in the custom ABR file format used by the Eaton Peabody Laboratories. Our CNN method outperforms EPL-ABR’s cross-correlation threshold estimation method across all metrics except for FPR on this dataset.

**Table 5:** Performance comparison of threshold estimation algorithms on Lab C data only (122 ABR waveforms from 2 mice). This table presents a side-by-side comparison of two threshold estimation algorithms: EPL-ABR’s cross-correlation algorithm and ABRA’s CNN-based thresholding algorithm. The metrics used for comparison include Accuracy, True Positive Rate (TPR), False Positive Rate (FPR), and the ability to estimate thresholds within 5,10, and 15 dB SPL. The values are presented as mean (± standard error). ABRA demonstrates superior or equal performance (green cells/bolded) in terms of accuracy, TPR, FPR, and estimating thresholds within 5, 10, or 15dB SPL.

| Metric \ Software | Cross-correlation (Suthakar & Liberman 2019) | ABRA Thresholding CNN |
| --- | --- | --- |
| Accuracy | 95.1 ($\pm 2.0$)% | **96.7 ($\pm 1.7$)%** |
| True Positive Rate | 95.6 ($\pm 2.2$)% | **97.8 ($\pm 1.6$)%** |
| False Positive Rate | **6.3 ($\pm 4.3$)%** | **6.3 ($\pm 4.3$)%** |
| Within 5 dB SPL | 91.7 ($\pm 8.0$)% | **100.00 ($\pm 0.0$)%** |
| Within 10 dB SPL | **100.00 ($\pm 0.0$)%** | **100.00 ($\pm 0.0$)%** |
| Within 15 dB SPL | **100.00 ($\pm 0.0$)%** | **100.00 ($\pm 0.0$)%** |

### Time Cost Analysis

In order to quantify the time savings of using the ABRA thresholding algorithm, a random sample of ABR files from 10 mice at 9 frequencies each for a total of 90 waveform stacks from Lab B was analyzed by two ABR raters from Lab A. It took both raters approximately 1 hour to manually analyze the ABR thresholds. However, using ABRA, it took about 48 seconds to output the automated thresholds for all frequencies, corresponding to 75x increased efficiency. The automated thresholds were within 5 dBs of Lab A inspection 90% of the time, 10 dBs 98% of the time, and 15 dBs 100% of the time. For comparison, inter-rater assessment showed that a Lab A annotator was within 5 dB of a Lab B annotator’s result 92% of the time, 10 dB 98% of the time, and 15 dB 100% of the time.

## Discussion

The deep learning techniques used in ABRA build on precedence not only in previous ABR studies but in the field of electrophysiology in general. A recent study showed that convolutional neural networks and long short-term memory architecture can automatically detect discrete records of protein movement (Celik et. al. 2020). Another electrophysiological study introduced a deep learning algorithm that infers latent dynamics from single-trial neural spiking data (Pandarinath et. al. 2018). This study used data from non-overlapping recording sessions that improved the inference of their model, similar to how our software accounts for a range of data collection protocols for improved generalizability. Both studies were designed to automate otherwise laborious and arduous tasks and simplify them such that future studies can be more accurate, more reproducible, and less time-consuming. The deep learning techniques used in our software have similar potential for ABR studies by streamlining the onerous task of labeling peaks and identifying thresholds. We envision future ABR acquisition protocols that can be guided by our software to avoid acquiring excess measurements after a threshold is reached. Together, this work sets the stage for massively accelerated electrophysiology experimental and analytical workflows, powered by deep learning-based algorithms.

While ABRA is a powerful tool set for ABR analysis, like all existing ABR analysis programs, it also has limitations that highlight areas for future development. While the CNN-based predictions of peak location are powerful, they offer little transparency. Future incorporation of interpretability methods may reveal waveform features that the model leverages to make accurate predictions across a large range of frequencies. Additional inclusion of biologically relevant priors in future versions may further reduce errors in peak finding. As for model limitations, the CNN-based thresholding model was trained only on mouse ABRs which had step sizes no larger than 20 dB; future work could extend or train new deep learning models to handle non-murine ABRs, which may have reduced signal-to-noise due to larger distances from the source generators. Validation of automated amplitude and latency measurements has so far been restricted primarily to wave 1, leaving waves 2-5 currently unvalidated; which can be pursued in future efforts as the model continues to incorporate new data from the labs mentioned here and others.

Another important consideration is the diversity of datasets used for model training: ABRA’s models have been trained on a diverse dataset including accelerated aging mouse models, mice with and without cadherin-23 correction, and two mouse lines exposed to varying noise exposures (**Supplementary Table 1**). However, this dataset is not exhaustive, and other datasets, especially those involving severe mutations, damage, or disease conditions may be significantly different from the training data. Such conditions, representing “out-of-distribution” cases, may require retraining or fine-tuning of existing models. To address this challenge, transfer learning methods could be employed, rapidly adapting existing deep learning models to new data conditions with minimal additional training data. Furthermore, ABRA’s modular design allows its deep learning models to be used independently from the GUI, facilitating integration into other researchers’ computational workflows and software environments. Most importantly, the accuracy of peak and threshold detection may not yet match that of the most seasoned experts in visual ABR analysis for abnormal, ambiguous, or low signal-to-noise waveforms. While the time saved by automation may still yet be a worthwhile tradeoff for certain applications, an additional benefit is the deterministic nature of the model and therefore high reproducibility. Most importantly, we anticipate significant improvements in performance as larger and more diverse datasets are incorporated over time.

ABRA has been designed to be a multi-purpose and versatile suite of tools and accompanying web app with extended functionality to be able to handle datasets acquired from different mouse strains and experimental settings (**Figure 9; Supplementary Information: The ABRA Graphical User Interface**). It includes readers to facilitate processing of datasets recorded in different formats, including the widely used standard .arf files from BioSigRZ Tucker Davis Technology recordings, .tsv/.asc files from EPL’s Cochlear Function Test Suite, or a generalized .csv file format from any number of other systems. ABRA’s automated thresholding method also reduces the time required for thresholding analyses by more than 50x compared to manual analysis and can streamline the process of extracting ABR thresholds from multiple subjects. In addition, the results can be exported to a .csv file for post-processing by the experimenter, and plots can be directly exported for publication if desired.

**Figure 9:**
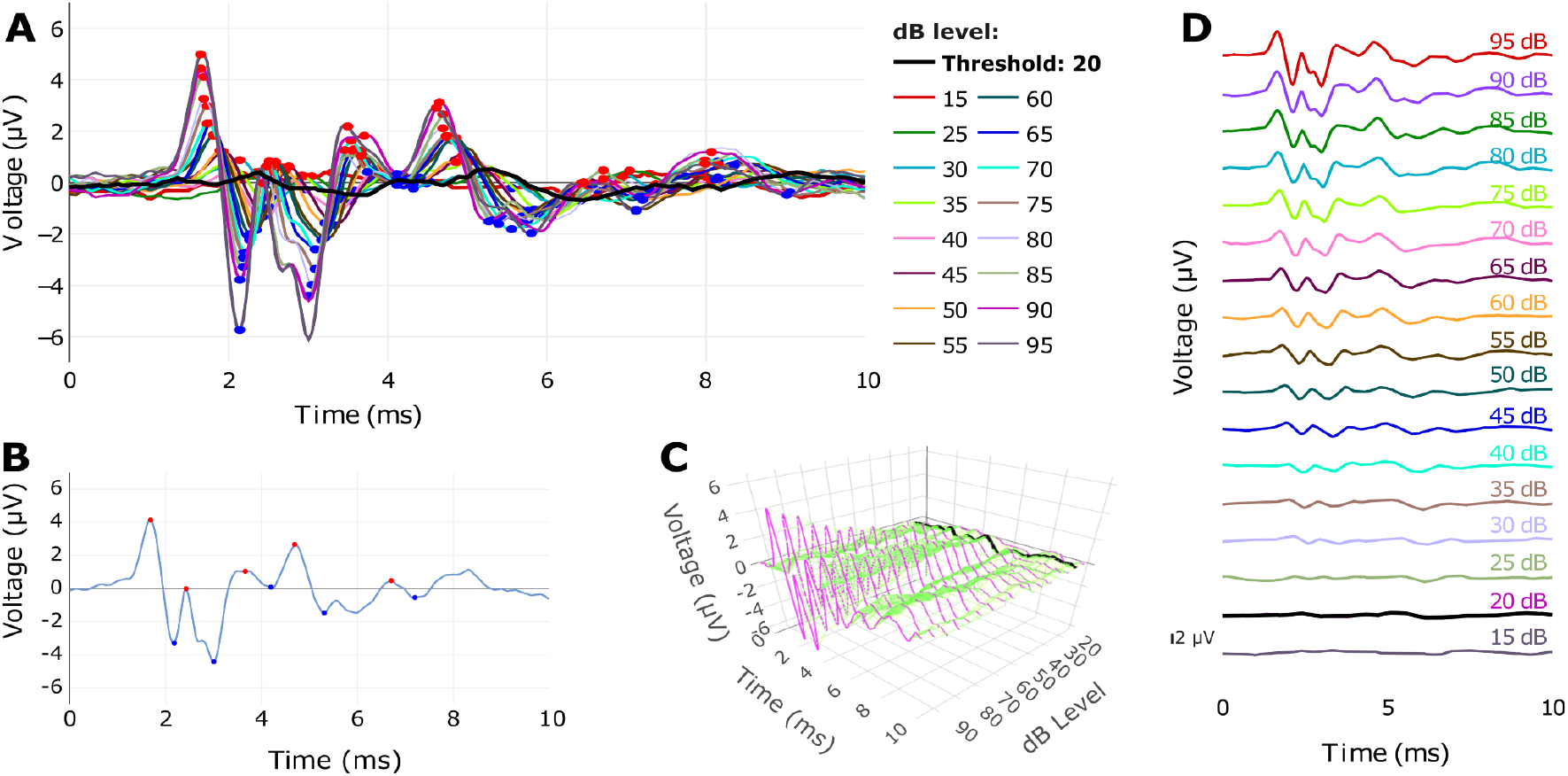
Visualizations of the combined ABRA thresholding and peak finding outputs. (A) Several ABR waveforms from a 1-month-old male C57Bl/6N mouse across sound amplitudes at 18 kHz, with predicted peaks and troughs (red and blue circles, respectively) and the predicted threshold (thick black line). (B) A single waveform (18 kHz; 85 dB SPL) with peaks and troughs labeled. (C) 3D representation of the same ABR waveforms in (A), with the predicted threshold (20 dB SPL) in black. (D) Stacked waveforms of the waveforms in (A), clearly showing the predicted threshold (20 dB SPL; black).

In summary, ABRA represents a significant advancement in auditory neuroscience, merging cutting-edge deep learning technology with accessible and intuitive software design. By automating the analysis of auditory brainstem responses—a crucial in vivo indicator of auditory function—ABRA not only accelerates hearing research but also enhances reproducibility and consistency across diverse experimental settings. The accompanying user-friendly graphical interface, coupled with powerful computational models, makes advanced data processing accessible to researchers from various backgrounds, including neuroscientists, audiologists, and computational biologists. Beyond hearing research, ABRA exemplifies how artificial intelligence can effectively tackle complex biological data analysis, offering valuable insights into sensory neuroscience, neurodegenerative diseases, and beyond. Ultimately, ABRA provides a versatile and scalable solution, poised to facilitate transformative discoveries at the intersection of biology, medicine, and computer science.

## Author contributions

A.E. and J.C. performed the majority of the model training and validation work and wrote the first draft of the manuscript. C.M.M. contributed to coding and validation work. E.C. assisted in generating training data and validating the model. S.B. helped with data annotation. Y.M.K. contributed to software development. R.A.F. provided funding support and expertise on ABR analysis. A.L. provided expertise on ABR analysis, assisted in validating the model, and helped interpret results. F.C. and W.M. contributed data and aided in interpreting results. C.C. provided expertise on statistical analysis and model development. C.C. and U.M. guided software development and analysis and co-supervised the overall project. U.M. conceived the project and provided funding support. All authors contributed to manuscript editing and reviewed the manuscript.

## Acknowledgements

We would like to thank Kirupa Suthakar and Charles Liberman for providing us with data as well as helping us improve our software, Zhijun Shen, Mark Rutherford, and Kali Burke for helping us improve our software and for insightful feedback on the manuscript, and Leah Ashebir and Peeyush Patel for help creating a manual for working with the ABRA GUI.

## Funding

U.M. is supported by a CZI Imaging Scientist Award (DOI:10.37921/694870itnyzk) from the Chan Zuckerberg Initiative DAF, NSF NeuroNex Award 2014862, the David F. And Margaret T. Grohne Family Foundation, the G. Harold & Leila Y. Mathers Foundation, NIDCD R018566-03S1 and R01 DC021075-01, and the UCSD Goeddel Family Technology Sandbox.

The collection of Marcotti Lab data was supported by an Innovation Seed Fund from the RNID (F115) to F.C. and BBSRC grant (BB/V006681/1) to W.M.

## Notes

https://abra.ucsd.edu

https://github.com/ucsdmanorlab/abranalysis

### Competing Interest Statement

The authors have declared no competing interest.

### Summary of Updates

In response to thoughtful suggestions, we have made the following key revisions: Refined Manuscript Focus: We have shifted the manuscript's emphasis from the software interface to the novel deep-learning algorithms themselves, moving details about the GUI to the supplementary information to better highlight the core scientific contribution. Strengthened Algorithm and Data Discussion: We have provided more detailed analyses of the datasets, including potential latency shifts between labs and considerations for aliasing during resampling. We also clari-fied the two-step peak-finding process to more accurately describe how the CNN's initial prediction is refined. Enhanced Data Visualization and Statistical Rigor: Several figures have been revised for improved clarity and accessibility, including using histograms and semi-transparent data points to better represent error distribu-tions. We have also incorporated DeLong's test for robust statistical comparison of model performance. Improved User Guidance and Functionality: Based on feedback regarding the software's usability, we have implemented more interactive features allowing users to manually adjust automatically-generated peaks and thresholds. We have also added warnings to the GUI to encourage critical evaluation of the results.

