## Supplementary Materials for "An Open-Source Deep Learning-Based Toolbox for Automated Auditory Brainstem Response Analyses (ABRA)"

### Supplementary Information

### Getting Started with ABRA

A tutorial manual for using the ABRA tools can be found on the tool's Github (https://github.com/ucsdmanorlab/abranalysis) and at the following link.

### Details on Data Collection

The multi-lab dataset used here (summarized in **Table 1**), was collected from three separate labs, with data collection details shown in **Supplementary Table S1** below.

| Methods | Lab A | Lab B | Lab C |
|---|---|---|---|
| **Anesthesia** | Ketamine (90 mg/kg) + Xylazine (10 mg/kg) | Ketamine (100 mg/kg) + Xylazine (10 mg/kg) | Ketamine (100 mg/kg) + Xylazine (10 mg/kg) |
| **Environment** | Soundproof chamber, heating pad (37°C) | Soundproof chamber, heating pad (37°C) | Soundproof chamber, heating pad (37°C) |
| **Electrode Placement** | Subcutaneous recording electrode at vertex, reference behind right pinna, ground on left leg | Subdermal electrodes behind pinna (reference and ground), vertex (active) | Needle electrodes: vertex to ipsilateral pinna (recording), ground near tail |
| **Sound Stimuli** | 5-ms tone pips (0.5 ms cos2 rise-fall), 21/sec | 5-ms pips (1.0-ms rise-fall with cos2 onset envelope), 42.6/sec | 5-ms pips (0.5-ms rise-fall with cos2 onset envelope), 30/sec |
| **Recording** | Filtered (300 Hz - 3 kHz), averaged using BioSigRZ software, 512 responses averaged | Customized software (Ingham et al., 2011), RZ6 auditory processor, 256 responses averaged | Amplified (10,000X), filtered (100 Hz - 3 kHz), averaged with A-D board in LabVIEW system, 1024 responses averaged |
| **Recording sampling** | 244 samples over 10 ms | 1953 samples over 20 ms | 426 samples over 17 ms |
| **Probed frequencies** | 4kHz, 8kHz, 16kHz, 24kHz, 32kHz | 100 Hz, 3kHz, 6kHz, 12kHz, 18kHz, 24kHz, 30 kHz, 36kHz, 42 kHz | 8kHz, 11.3kHz, 16kHz, 22.6kHz, 32kHz, 45.2kHz |
| **Sound Intensity** | Decreased from 90 dB SPL to 10/20 dB SPL in 5 dB steps | 0-95 dB SPL in 5 dB steps | Raised from ~10 dB SPL below threshold to 80 dB SPL in 5 dB steps |
| **Speaker distance** | Open-field - 10 cm from ear | Open-field - 10 cm from ear | Closed-field - ~3 cm from the eardrum |
| **Mouse age/strains used** | 3-month SAMP8 (Senescence-Accelerated Mouse-Prone 8) (Takeda et al. 1981) | 1-month C57Bl/6N with and without corrected CDH23 | 7-week C57Bl/6J and CBA/CaJ, after varying noise exposures (Wu et al. 2024) |

**Supplementary Table S1**: **Summary of the experimental recording conditions used by the three labs.** The specific methods employed by each lab—Manor Lab (Lab A), Marcotti Lab (Lab B), Liberman Lab (Lab C) in collecting each dataset are summarized, including anesthesia, environment, electrode placement, sound stimuli, response recording, sound frequencies and intensity, distance between the mouse and speaker, and mouse strains and ages.

**The ABRA Graphical User Interface**

The ABRA GUI incorporates the ABRA peak finding and thresholding tools, and was developed in Python using the Streamlit framework ("Streamlit" n.d.), providing an interactive platform for researchers to visualize ABR data. All documentation of the code for the graphical user interface (GUI) and instructions for using the ABRA tools can be found at: https://doi.org/10.5281/zenodo.15054979 (Erra et al. 2025). The ABRA GUI allows users to import multiple ABR data files, and accepts data in most commonly used formats: in .arf or exported .csv formats for data collected with the BioSigRZ software (Tucker Davis Technologies) and in .asc or .tsv format for data collected with the CFTS software (Eaton-Peabody Laboratories). For other data types, the application will accept any data converted into a generalized .csv format, and an example template is provided. Upon import, the data is preprocessed to extract the timescale of the recording, the stimulus frequencies and amplitudes, and the waveform data.

After import and preprocessing, the GUI allows the user to select which frequencies and decibel levels they wish to examine for plotting and analysis. The ABR plots are shown through the Plotly framework in Python and can be downloaded as .png and .pdf files (Plotly Technologies Inc. 2015). Calculated metrics related to the displayed waveforms are displayed under the plots, including wave 1 amplitude, latency to the first peak, and threshold (as defined in **Figure 1**). These metrics can be downloaded as a .csv file. The plotting functions allow the user to view all the waveforms for a single frequency, highlight the automatically detected peaks and troughs, and automate thresholding (**Figure 9**).

The ABRA interface also implements two novel visualization features for ABR waveforms of varying stimulus amplitudes at a given stimulation frequency. First, it provides the option to implement time warping, which visually aligns the peaks and troughs of multiple waveforms (see **Supplementary Figure S1**). This view does not change the underlying data, and does not affect wave amplitudes, but stretches and compresses the waveform to varying degrees along the time axis to enhance visualization. Second, the app provides an interactive 3D surface plot of waveforms which allows the user to view the series of ABR waveforms as a surface in the 3-dimensional space created by the time domain, the probed decibel levels, and the recorded ABR voltage. These various functionalities can provide the user with tools to visually ascertain features like thresholds and peaks and automate model predictions of those features in tabular form. The ABRA tool set also provides automated batch analyses for multiple data files.

**ABR Curve Alignment with Time Warping**

ABRs from mice exhibit a characteristic structure with 5 distinct peaks (**Figure 1**). However, a common challenge in analyzing these ABR waveforms is the non-uniform latency across different frequencies and decibel levels. This variability in latency can distort functional summary statistics (e.g. mean ABR curve, covariance surface) and time-based comparisons of these responses, as the peaks do not occur at the same time across different ABRs for the same mouse. To address this, we provide an option to employ time warping to align these ABRs, which aligns the position of peaks and other salient features of the ABRs across time. This alignment decouples amplitude from latency variation, facilitating the visual comparison of amplitudes of ABR waveforms. The encoding of time alignment parameters into individual-specific warping functions provides the option of incorporating these features into machine learning models, which in some cases improves the models' performance and predictive power as it did for the Logistic Regression and XGBoost Classifiers for automated thresholding. Because time warping adjusts the spacing between points on the time axis (without affecting the amplitude), the time warped curves can be used to visually inspect threshold or peak amplitude, but not latency, which should be assessed on the original unwarped curves.

To conduct the time warping step, we use the *fdasrsf* package in Python (Tucker 2021). This package implements elastic time warping, a method that maximizes the alignment of key features in waveforms. Here, this technique provides aligned ABRs, counteracting the non-uniform latency across different frequencies and decibel levels.

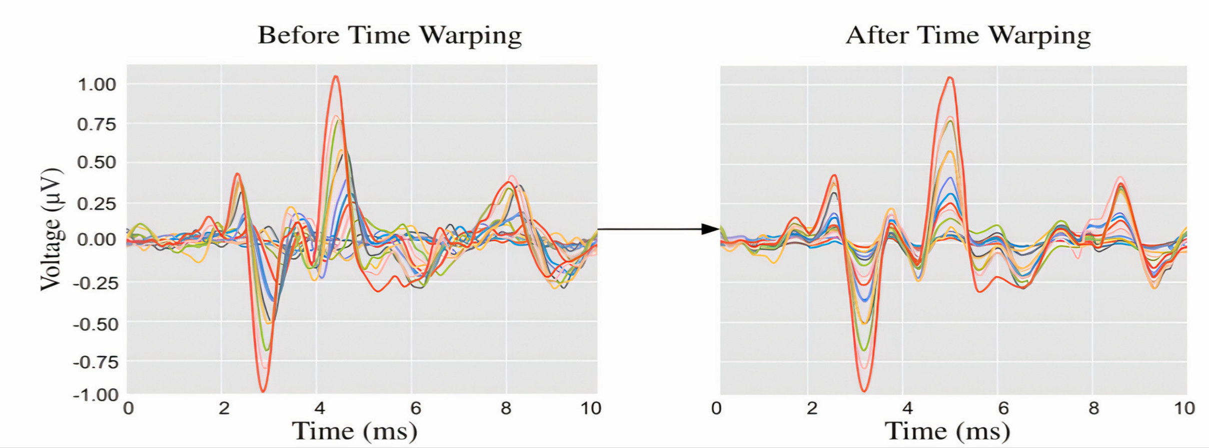


**Supplementary Figure S1: ABRs before (left) and after (right) Time Warping.** The depicted transformation of waveforms, both before and after applying elastic time warping using the *fdasrsf* package (Tucker 2021), illustrates clear registration of waveform features. Associated with each waveform is also an estimated time warping function which is useful in quantifying changes between the original unaligned latencies and the aligned latencies for all wave peaks and troughs.

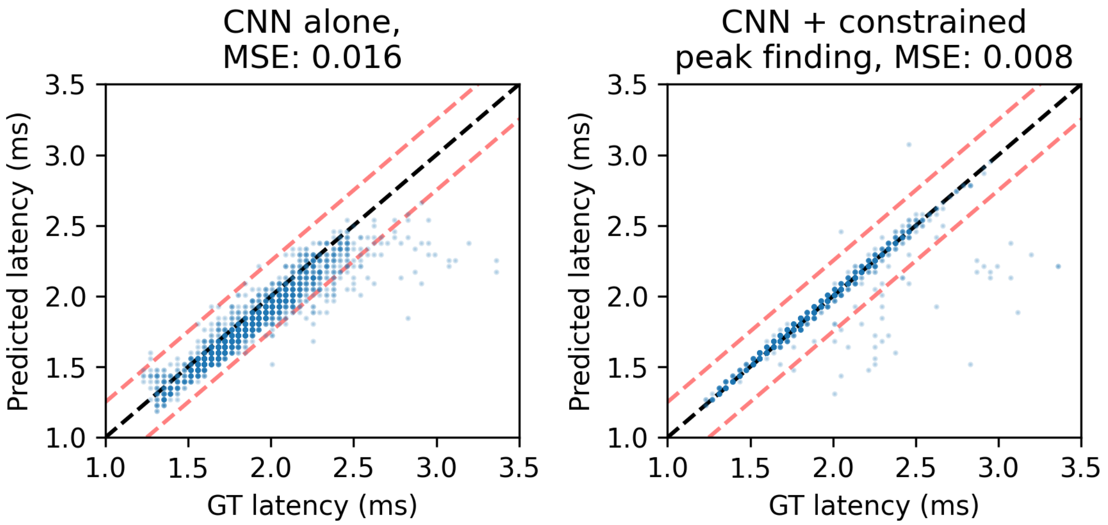


**Supplementary Figure S2: Peak 1 detection, CNN alone vs. CNN with constrained peak finding.** Predicted vs ground truth (GT) latency is shown for the direct output of the CNN (left) and the CNN output followed by the constrained peak finding function (right). Red dashed lines show +/- 0.25 ms error margins.

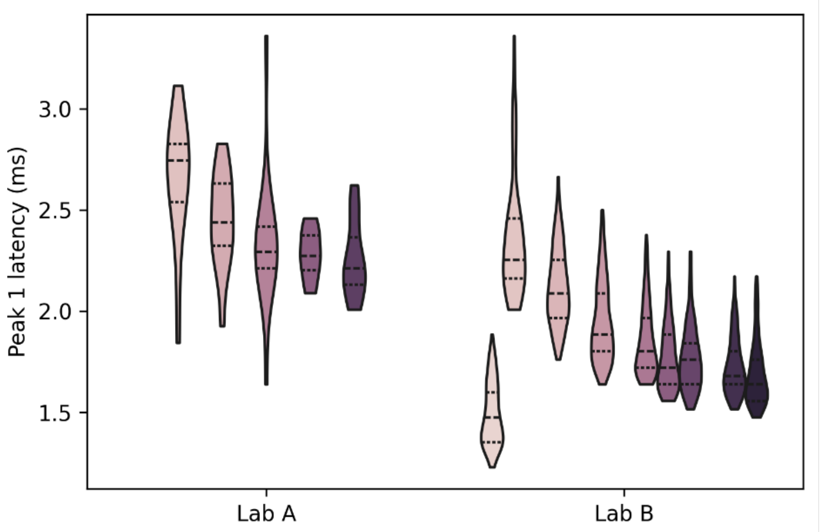


**Supplementary Figure S3: Ground truth annotations for peak 1 latency.** Annotations from Lab A include responses to 4, 8, 16, 24, and 32 kHz sound (left to right), and from Lab B include responses to Click, 3, 6, 12, 18, 24, 30, 36, 42 kHz sound (left to right).

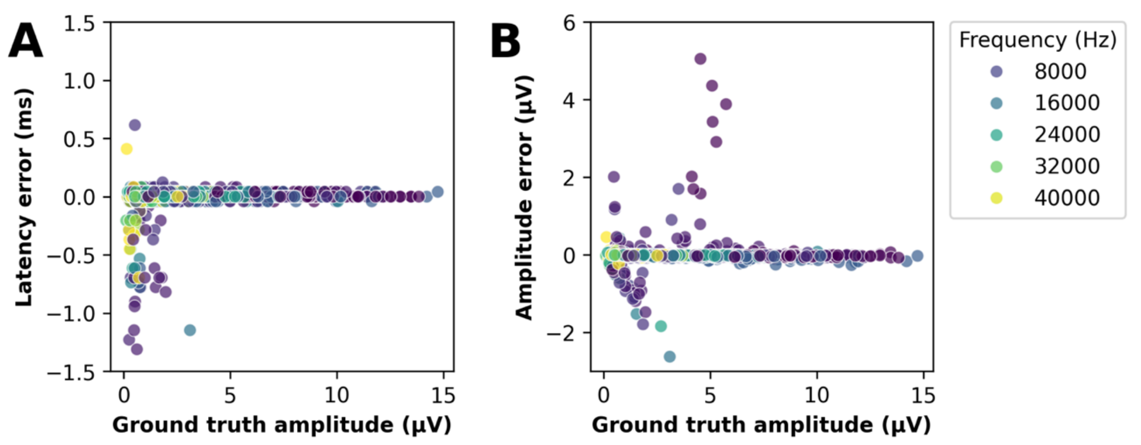


**Supplementary Figure S4: Peak 1 errors by amplitude and frequency.** Wave 1 latency and amplitude errors vs. ground truth wave 1 amplitude, as in Figure 5C, 5D. Each point is a single prediction, color-coded by frequency.

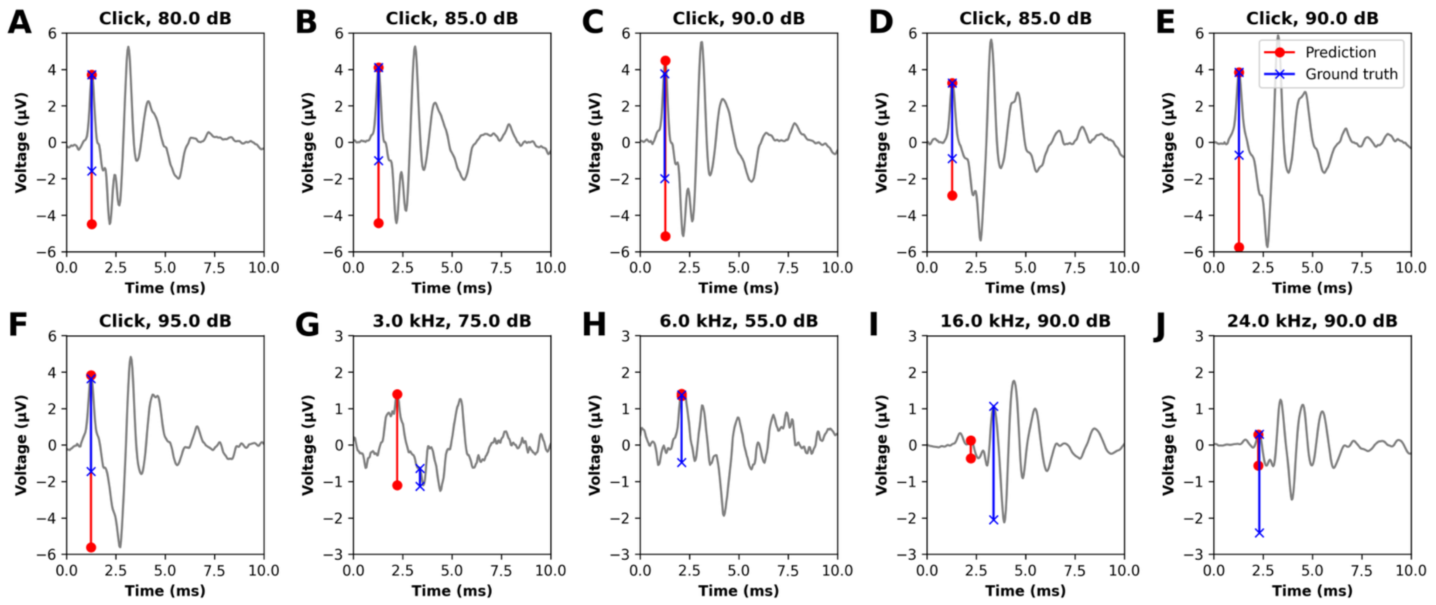


**Supplementary Figure S5: The ten highest magnitude peak 1 amplitude errors.** For each waveform, the predicted (red circles) and "ground truth" annotations (blue x's) are shown. The top point lies at the peak, and the line dropping down represents the peak amplitude. These top 10 errors come from a small number of cases: A, B, and C are Click responses from one mouse; D, E, and F are Click responses from a second mouse. H is an erroneously small amplitude due to a small trough near the top of peak 1. I and J are annotator errors.

### ML Model Hyperparameters

Optimal choices of hyperparameters chosen by cross-validation for each of the thresholding and peak-finding models are displayed in Table S2 below.

| | Thresholding CNN | Thresholding XGB | Thresholding LR | Peak Finding CNN |
|---|---|---|---|---|
| Hyper-parameters | Loss function: Binary Cross Entropy<br>Activation function for all layers except final layer: Relu<br>Final activation function: Sigmoid<br>Batch Size: 128 | Subsample size for each tree: 0.8<br>Positive class weight: 2<br>Boosting rounds: 700<br>Minimum sum of instance weight (hessian) in a leaf node: 1 | All parameters are Scikit-learn defaults | Loss function: Mean Squared Error<br>Activation function for all layers: Relu<br>Batch Size: 32<br>Early Stopping Patience: 25<br>Optimizer: Adam |

| | | | | |
|---|---|---|---|---|
| | Early Stopping Patience: 25<br>Reduce Learning Rate on Plateau Patience: 20<br>Optimizer: Adam<br>Learning Rate: 1e-4<br>Conv. Layer 1 Filters: 128<br>Conv. Layer 2 Filters: 128<br>Conv. Layer 3 Filters: 64<br>Conv. Layer Stride: 1<br>Conv. Layer Padding: 0<br>Kernel Size: 7<br>MaxPool Size: 2<br>MaxPool Stride: 2<br>MaxPool Padding: 0<br>Fully Connected Layer Size: 128<br>Dropout Rate between Conv. Layers and before first Fully Connected Layer: 0.5<br>Dropout Rate between Fully Connected Layers: 0.4 | Maximum depth of a tree: 5<br>Learning Rate: 0.05<br>Gamma (Regularization parameter for tree splitting): 0.5<br>Fraction of features used for fitting each tree: 1.0 | | Learning Rate: 1e-3<br>Weight Decay: 1e-5<br>Conv. Layer 1 Filters: 128<br>Conv. Layer 2 Filters: 32<br>Conv. Layer Stride: 1<br>Conv. Layer Padding: 1<br>Kernel Size: 3<br>MaxPool Size: 2<br>MaxPool Stride: 2<br>MaxPool Padding: 0<br>Fully Connected Layer Size: 128<br>Dropout Rate between Conv. Layer 1 and Conv. Layer 2: 0.5<br>Dropout Rate between Conv. Layer 2 and first Fully Connected Layer: 0.3<br>Dropout Rate between Fully Connected Layers: 0.1 |
| Library | Keras (3.3.3) | XGBoost (1.7.3) | Scikit-learn (1.2.2) | Pytorch (2.2.0.post100) |

**Supplementary Table S2: Candidate Model Hyperparameters for Thresholding and Peak Detection Models.**
